# Clinically relevant mutations of mycobacterial GatCAB inform regulation of translational fidelity

**DOI:** 10.1101/2021.03.29.437598

**Authors:** Yang-Yang Li, Rong-Jun Cai, Jia-Ying Yang, Tamara L. Hendrickson, Ye Xiang, Babak Javid

**Author notes:** These authors contributed equally (Yang-Yang Li, Rong-Jun Cai).

## Abstract

Most bacteria employ a two-step indirect tRNA aminoacylation pathway for the synthesis of aminoacylated tRNA^Gln^ and tRNA^Asn^. The heterotrimeric enzyme GatCAB performs a critical amidotransferase reaction in the second step of this pathway. We have previously demonstrated in mycobacteria that this two-step pathway is error-prone and translational errors contribute to adaptive phenotypes such as antibiotic tolerance. Furthermore, we identified clinical isolates of the globally important pathogen *Mycobacterium tuberculosis* with partial loss-of-function mutations in *gatA*, and demonstrated that these mutations result in high, specific rates of translational error and increased rifampicin tolerance. However, the mechanisms by which these clinically-derived mutations in *gatA* impact GatCAB function was unknown. Here, we describe biochemical and biophysical characterization of *M. tuberculosis* GatCAB, containing either wild-type *gatA* or one of two *gatA* mutants from clinical strains. We show that these mutations have minimal impact on enzymatic activity of GatCAB; however, they result in destabilization of the GatCAB complex as well as that of the ternary asparaginyl-transamidosome. Stabilizing complex formation with the solute trehalose increases specific translational fidelity of not only the mutant strains, but also of wild-type mycobacteria. Therefore, our data suggest that alteration of GatCAB stability may be a mechanism for modulation of translational fidelity.

## Introduction

Protein synthesis is an essential, but error-prone biological process (1-3). Recent evidence suggests that optimal translational fidelity is not necessarily as high as possible. This consequence is not just due to reasons of efficiency, i.e. a trade-off between speed and accuracy of protein synthesis, but also because translational errors, mistranslation, may allow adaptation to hostile environments, particularly in contexts of environmental stress (3-5). Nonetheless, excessive mistranslation is harmful. Therefore, mechanisms for ‘tuning’ of optimal – and context-specific – translational fidelity are required. Multiple mechanisms for microbial adaptive mistranslation have been proposed (2,3). Most bacterial species, including the majority of pathogens use an indirect tRNA aminoacylation pathway due to their lack of specific glutaminyl- or asparaginyl tRNA synthetases – or both (6). In mycobacteria, which lack both glutaminyl- and asparaginyl-synthetases, we have previously shown that baseline error rates associated with the pathway (i.e. glutamine to glutamate and asparagine to aspartate) are ∼0.2-1% /codon, orders of magnitude higher than equivalent errors in *Escherichia coli*, which lacks the pathway (7,8). Furthermore, elevated mistranslation in this pathway causes increased tolerance to the first-line anti-mycobacterial drug rifampicin due to gain-of-function protein variants arising from mistranslation of critical residues in the drug target of rifampicin, the beta subunit of RNA polymerase (7,9). Consistently, reducing errors in this pathway with the small molecule kasugamycin increases susceptibility to rifampicin in *M. tuberculosis*, both in axenic culture and in animal infection (10), confirming the critical role for mistranslation in mycobacterial rifampicin tolerance.

In the first step of the two-step indirect pathway, a non-discriminatory glutamyl-(ND-GluRS) or aspartyl-tRNA synthetase (ND-AspRS) mischarges tRNA^Gln^ to form Glu-tRNA^Gln^ and tRNA^Asn^ to form Asp-tRNA^Asn^ respectively (11,12). These mischarged tRNAs are specifically recognized by the heterotrimeric amidotransferase GatCAB, which converts them to Gln-tRNA^Gln^ and Asn-tRNA^Asn^ respectively. Mutations in *gatCAB* are surprisingly common, and have been identified in a substantial minority of clinical isolates of *M. tuberculosis* sequenced genomes; several of these mutations cause both elevated rates of mistranslation and rifampicin antibiotic tolerance (9). This is all the more remarkable since these mutants were in disease-causing strains isolated from patient samples, suggesting that mycobacteria are surprisingly plastic and tolerant to elevated translational error arising from this pathway.

Bacterial GatCAB is a heterotrimeric, glutamine-dependent enzyme complex composed of GatA, GatB and GatC. Based on primary sequence analysis, reported crystal structures and experimental analyses of bacterial GatCAB (13-15), GatA serves as the glutaminase subunit for ammonia production (11), GatB is responsible for activation and transamidation of the misacylated tRNA (16), and GatC is a small scaffold protein that helps stabilize the complex (13). In some bacteria, GatCAB forms a ternary complex with ND-AspRS and possibly with ND-GluRS, and tRNA^Asn^ or tRNA^Gln^ respectively, termed the transamidosome (17-22). It has been proposed that due to the physical proximity of the non-discriminatory synthetase (the source of potential translational error) and GatCAB (which corrects mischarged tRNA complexes), the transamidosome may promote the efficiency of and minimize errors due to this pathway. Here we biochemically and biophysically analyse mycobacterial GatCAB as well as two mutant enzymes. We show that the mutations primarily function to destabilise the enzyme and the ternary transamidosome complex. Stabilization of the complex with trehalose decreases mistranslation rates in both mutant and wild-type mycobacteria, suggesting that enzyme and transamidosome stability are mechanisms for regulating translational fidelity in mycobacteria.

## Material and Methods

### Bacterial strains and growth conditions

Details of constructed strains and primers used in this study are listed in Supplementary Tables S1 and S2. *E. coli* strains were cultured in LB medium (BD Difco, #DF0402-07-0) or 2×YT medium with antibiotics as appropriate. All *Mycobacterium smegmatis* strains were cultured in Middlebrook 7H9 medium (BD Difco, #DF0713-17-9) supplemented with 0.2% glycerol, 0.05% Tween-80, 10% ADS (albumin-dextrose-salt), in the presence of hygromycin (50 µg/mL) or trehalose, as indicated. If not otherwise noted, cells were grown and maintained at 37°C with shaking.

### Cloning, expression and purification of *M. tuberculosis* ND-AspRS

The *M. tuberculosis aspS* gene (encoding ND-AspRS, start codon changed to ATG) containing 6×His-tag fused at the C-terminus, was amplified by PCR with the appropriate primers (Supplementary Table S2). The PCR product was digested with *Xba*I and *Hind*III, and inserted into the corresponding sites of pET-28a(+) (Novagen, #69864-3CN). The expression vector was transformed into *E. coli* Transetta (DE3) chemically competent cells (Transgen Biotech, #CD801). Cells were grown at 37 °C in LB medium containing 50 µg/mL kanamycin and 34 µg/mL chloramphenicol, and protein expression was induced by adding IPTG to a final concentration of 1 mM at an OD_600nm_ of 0.4-0.6 for 4 h. Cells were harvested, resuspended in buffer A (50 mM NaH_2_PO_4_ pH 7.4, 300 mM NaCl) containing 10 mM imidazole, 0.1 U/mL DNase I (Thermo Scientific, #EN0521) and InStab Protease Cocktail, EDTA-free (Yeasen Biotech, #20123ES50). After sonication and centrifugation at 11,000 rpm (15,557 *g*) for 75 min at 4 °C, the supernatant was loaded onto the column with pre-equilibrated Ni-NTA resin (Qiagen, #30230). All the following purification processes were carried out at room temperature, with buffers containing protease inhibitor stored on ice. The column was washed with 10 mM and 40 mM imidazole in buffer A sequentially, and the protein was eluted in a step-wise manner with 80mM, 150mM and 250 mM imidazole in buffer A respectively. Only the 250mM elution fractions were collected and exchanged into buffer B (50 mM NaH_2_PO_4_ pH 7.4, 300 mM KCl), and concentrated using Amicon Ultra-15 30 kDa centrifugal filter device (Merck Millipore, #UFC903024) at 4 °C. The final protein concentration was determined by Bradford reagent (Bio-Rad, #5000205) and the purity checked by SDS-PAGE. For aminoacylation assay, the protein was stored at −20 °C in buffer B containing 50% glycerol. Otherwise, ND-AspRS was flash frozen in liquid nitrogen and stored at −80 °C.

### Cloning, expression and purification of *M. tuberculosis* GatCAB

The *M. tuberculosis gatCA* gene (start codon of *gatC* and *gatA* changed to ATG) fused to an N-terminal Strep-tag II (amino acid sequence WSHPQFEK), and *gatB* gene containing a C-terminal 6×His-tag, were amplified by PCR with the appropriate primers (Supplementary Table S2), and ligated to *Nco*I/*Pac*I digested pETDuet-1 vector (Novagen, #71146-3CN), using Gibson Assembly (New England Biolabs, #E2611). The expression vector was transformed into *E. coli* Transetta (DE3) chemically competent cells. Cells were grown at 37 °C in LB medium containing 100 µg/mL ampicillin and 34 µg/mL chloramphenicol. Protein expression was induced by adding IPTG to a final concentration of 0.2 mM at an OD_600nm_ of 0.6-0.7, and bacteria cultured for 16 h at 16 °C. Buffers for all the following steps were added with protease inhibitor.

Cells were harvested, resuspended, sonicated and centrifuged in the same manner as ND-AspRS purification. The protein complex was first purified by Ni-NTA affinity chromatography, carried out at room temperature with buffers stored on ice. Clarified supernatant was bound to the column pre-equilibrated with buffer A containing 10 mM imidazole. After subsequent washing with 10 mM and 40 mM imidazole in buffer A, the proteins were eluted with buffer A containing 250 mM imidazole. The elution then went through buffer exchange for further purification, carried out at 4 °C.

For the GatA glutaminase assay, the Ni-NTA eluate was exchanged into buffer C (50 mM Tris-HCl pH 8.0, 300 mM NaCl), concentrated, and applied to Strep-Tactin affinity chromatography column (IBA Lifesciences, #2-1201-025) according to the manufacturer’s protocol. The protein was further purified by size-exclusion chromatography using ÄKTA purifier (Cytiva) and a HK 16/40 column (HUIYAN Bio, #HT16-40) packed with Sephacryl S-300 HR resin (Cytiva, #17-0599-10), equilibrated with buffer C. The elution buffer was buffer C containing 5 mM d-Desthiobiotin (Sigma-Aldrich, #D1411). Peak fractions eluted at the expected volume were pooled and concentrated. For other studies, the Ni-NTA eluate was exchanged into buffer D (20 mM HEPES-NaOH pH 8.0, 300 mM NaCl), concentrated, and applied to Strep-Tactin affinity chromatography according to the manufacturer’s protocol (IBA Life Sciences). The protein was eluted with buffer D containing 5 mM d-Desthiobiotin, exchanged into buffer D and concentrated.

The final protein concentration was determined by Bradford reagent or Qubit protein assay kit (Invitrogen, #Q33212), and the presence of all three subunits was verified by SDS-PAGE. For the GatCAB amidotransferase assay, the protein was stored at −20 °C in buffer D containing 50% glycerol. For other studies, GatCAB was flash frozen in liquid nitrogen and stored at −80 °C. GatCAB variants were generated using site-directed mutagenesis using appropriate primer pairs, and purified to homogeneity as above.

### Preparation of *M. tuberculosis* tRNA^Asn^

The T7 promoter and the Mtb tRNA^Asn^ gene were amplified using PCR with the appropriate primers (Supplementary Table S2). The PCR product was digested with *EcoR*I and *BamH*I and inserted into the corresponding sites of pTrc99a (TIANDZ, #60908-6580). The expression vector was transformed into *E. coli* BL21 (DE3) chemically competent cells (CWBio, #CW0809). Cells were grown at 37 °C in 2×YT medium containing 100 µg/mL ampicillin to OD_600nm_ of 0.8 before induction. Overexpression was induced by adding IPTG to a final concentration of 0.4 mM, and bacteria cultured for 16 h at 37 °C. Total nucleic acids were extracted with minor modifications of a previously described procedure (23). Cells were harvested and resuspended in 50 mM Tris-acetate pH 7.8, 4 M guanidine thiocyanate, 15 mM β-mercaptoethanol and 2% Triton X-100. After 15 min of incubation on ice, the mixture was mixed with an equal volume of 3M sodium acetate pH 6.5, incubated for another 15 min on ice, and centrifuged at 11,000 rpm (15,557 *g*) for 30 min at 4 °C. The lysate was precipitated with isopropanol followed by centrifugation at 11,000 rpm (15,557 *g*) for 10 min at 4 °C. The pellet was resuspended in 10 mM Tris-acetate pH 7.8 and 1 mM EDTA, and the suspension was extracted twice with equal volumes of RNA extraction reagent (Phenol:Choloform:Isoamyl alcohol = 25:24:1, pH < 5.0, Solarbio, #P1011). The aqueous layer was isopropanol precipitated. The pellet was re-dissolved in DEPC-treated water and incubated with 3 times volume of 100 mM Tris-HCl pH 9.0 for 60 min at 37 °C to ensure complete deacylation of the tRNA.

The mixture was isopropanol precipitated for subsequent tRNA enrichment based on the procedure of Spears *et al*. (24) Briefly, the total nucleic acid pool was mixed with 1 M MOPS pH 7.0 to a final concentration of 0.1 M, and loaded onto a Q-2500 column (Qiagen #10083) pre-equilibrated with 2× 50 mL of fresh buffer E (50 mM MOPS pH 7.0, 15% isopropanol, 1% Triton X-100). After washing the column with 4x 50 mL of buffer F (50 mM MOPS pH 7.0) containing 200 mM NaCl, the RNA was eluted with multiple fractions of 10 mL buffer F containing 650 mM NaCl. Ideal fractions of eluate (based on denaturing urea-PAGE, not shown) were collected and precipitated with isopropanol, and pellets were stored at −20 °C. This approach yielded a highly enriched stock of Mtb tRNA^Asn^ containing also *E. coli* tRNAs and other nucleic acids.

### Folding, quantification and aminoacylation of tRNA^Asn^

The experiment was performed as previously described (23) with some modifications. Prior to use for aminoacylation with aspartate, tRNA^Asn^ was dissolved in DEPC-treated water to the desired concentration, and incubated in a 75 °C water bath for 5 min. MgCl_2_ was added to a final concentration of 2 mM once the tRNA^Asn^ was allowed to cool slowly to 65 °C. After the sample was cooled to below 45 °C, it could be used for aminoacylation or stored at −20 °C for later experiments.

The fraction of chargeable tRNA^Asn^ was confirmed for every batch prepared. Briefly, an aminoacylation assay with ND-AspRS was performed as follows: 100 µL aminoacylation assay with 20 mM HEPES-OH pH 7.5, 2 mM ATP, 4 mM MgCl_2_, 100 µM aspartate, and 25 µCi/mL L-[2,3-^3^H] aspartate (PerkinElmer, # NET390V001MC). The reaction was initiated by adding 1 µM ND-AspRS followed by incubation at 37 °C for 1 h. The assay was quenched by adding phenol/chloroform (pH < 5.0). The aqueous layer was isopropanol-precipitated at −20 °C for 2 hours and then pelleted. The pellet was dissolved in 100 µL ddH_2_O, from which aliquots of 10 μL were counted in vials with 3 mL OptiPhase HiSafe3 scintillation fluid (PerkinElmer, #1200-437). The fraction of charged tRNA^Asn^ was calculated based on the charged tritium labeled aspartate incorporated into the pellet. Unlabelled Asp-tRNA^Asn^ was prepared as above but without any tritium-labelled aspartate. The pelleted Asp-tRNA^Asn^ was dissolved in DEPC-treated water to 200 μM and stored in small aliquots at −20 °C if not used immediately.

### GatA glutaminase assay

The GatA glutaminase assay was conducted at 37 °C in 40 mM HEPES-KOH pH 7.5, 25 mM KCl, 8 mM MgCl_2_, 5 µM Asp-tRNA^Asn^. The concentration of GatCAB was ∼20 nM. The reaction mixture for each assay was incubated on ice then pre-equilibrated at 37 °C for 2min. Glutamine (final concentration varying from 50-2000 µM) was added to initiate the reaction. At each time point, a 20 µL aliquot was removed and quenched with 10 µL 0.3 M HCl, followed by the addition of 10 µL 450 mM Tris-HCl pH 8.0. The amount of glutamate in the mixture was measured using the Glutamate-Glo Assay (Promega, #J7022) according to the manufacturer’s instructions.

### Labeling and aminoacylation of ^32^P-labeled *M. tuberculosis* tRNA^Asn^

The experiment was performed as previously described (23) with some modifications. *M. tuberculosis* tRNA^Asn^ (final concentration 2.5 µM) was radiolabeled at 37 °C for 30 min with 1 µM *E. coli* tRNA nucleotidyltransferase and [α-^32^P] ATP (final activity 2 µCi/µL) (PerkinElmer, # BLU003X250UC) in 50 mM Tris-HCl pH 8.0, 20 mM MgCl_2_, 5 mM dithiothreitol (DTT), and 50 µM sodium pyrophosphate (NaPPi). After phenol/chloroform (pH < 5.0) extraction, free [α-^32^P] ATP was removed from the sample via a MicroSpin G-25 column (Cytiva, #27-5325-01) and then the tRNA was isopropanol-precipitated at −20 °C for 2 h. ^32^P-labeled tRNA^Asn^ (0.4 µM) was diluted with 12.5 µM unlabeled tRNA^Asn^ and this diluted mixture was used for aminoacylation assays as described above using unlabeled aspartate. The ^32^P-labeled Asp-tRNA^Asn^ pellet was dissolved in ddH_2_O to 80 µM and stored in aliquots at −20 °C until needed.

### ^32^P nuclease P1 GatCAB amidotransferase assay

The experiment was performed as previously described (23) with some modifications. The assay was performed in 40 mM HEPES-KOH pH 7.5, 25 mM KCl, 8 mM MgCl_2_, 4 mM ATP, 1 mM glutamine, 10 nM GatCAB and ^32^P-labeled Asp-tRNA^Asn^ (various concentration from 0.25-8.0 µM). The reaction was kept on ice until the 2 min pre-equilibration step at 37 °C, and then initiated with ATP and glutamine. A 5 µL aliquot for each time point was removed and quenched with 5 µL nuclease P1 (Sigma #N8630) in a 100 mM sodium citrate suspension (pH 4.7, 0.66 mg/mL). The digestion was kept in a 37 °C heat block for 30 min, and 3 µL of sample was carefully spotted onto pre-washed PEI-cellulose TLC plates (Merck Millipore, #105725). The TLC plate was eluted in developing buffer containing 10 mM NH_4_Cl and 5% acetic acid for 100-140 min. The air-dried plates were exposed to phosphor screens for at least 16 h, and then quantified by Typhoon FLA 9500 phosphorimager (Cytiva).

### Gel filtration analysis

Experiments were carried out at 4 °C using an ÄKTA purifier and Superdex 200 Increase 10/300 GL column (Cytiva, #28-9909-44). 500 µL samples with indicated component concentrations were prepared in buffer G (50 mM HEPES-KOH pH 7.2, 30 mM KCl, 6 mM MgCl_2_, 1 mM DTT), incubated on ice for 30 min, and loaded onto the column equilibrated with buffer G. Peak fractions were pooled, concentrated using an Amicon Ultra-0.5 3 kDa centrifugal filter device (Merck Millipore, #UFC500324) at 4 °C, and characterized by SDS-PAGE and native PAGE. For native PAGE analysis, an 8% polyacrylamide gel was prepared, run in buffer H (buffer G without 1 mM DTT) for 2 h at 90 V on ice, and stained with SYBR Green II RNA gel stain (Solarbio, #SY1040). A Qubit protein assay kit was used to measure the protein concentration of the sample, which was then diluted to 0.5 mg/mL for the stability assays.

### *In vitro* stability assay

The UNcle instrument (Unchained Labs) was used to measure the melting and aggregation of samples over a thermal ramp. 9 µL of 1.7 mg/mL GatCAB in buffer D or 0.5 mg/mL transamidosome in buffer G were loaded into a Uni (sample holder for 16 quartz capillary tubes used for analysis). Buffers were supplemented with 0.2 / 0.4 M trehalose as indicated. The temperature increased from 20 to 90 °C, in constant increments of 0.3 °C/min, and the instrument generated both the BCM (barycentric mean of the intrinsic fluorescence spectra versus temperature) and aggregation curves (static light scattering intensity using lasers at 266 nm versus temperature). To determine Tm1, namely the mid-point temperature of the first unfolding transition, the first-order derivative of the BCM curve was calculated and plotted against temperature to generate the derivative curve; the temperature corresponding to the first peak of the derivative curve represents Tm1. The aggregation onset temperature (Tagg_266_), was calculated by the UNcle Analysis software, with the Tagg_266_ peak threshold value set as 30%.

### Structural modelling of *Mycobacterium tuberculosis* Asn-transamidosome

A model of the *M. tuberculosis* Asn-transamidosome was constructed based on the structure of the *Pseudomonas aeruginosa* transamidosome (PDB ID:4WJ3) (20). GatCAB and ND-AspRS sequences were from *Mycobacterium tuberculosis-*H37Rv, and the structures were built using SWISS homology modelling (https://swissmodel.expasy.org). The tRNA^Asn^ sequence was from *P. aeruginosa*.

### *In vivo* mistranslation assay

Mistranslation rates were measured in strains with *M. tuberculosis gatCA*-WT, G444S and K61N, constructed on an isogenic *M. smegmatis gatCA* deletion background with *Renilla*-Firefly dual-luciferase reporters (9). The construct containing D120N in the *Renilla* luciferase sequence was used to measure the asparagine-to-aspartate mistranslation rate, while the construct containing K529R in the Firefly luciferase sequence was used to measure the near-synonymous arginine-to-lysine mistranslation rate. The assay was carried out as previously described (7,25,26). Briefly, strains containing mistranslation reporters were grown to stationary phase (OD_600nm_ > 3) and diluted into fresh 7H9 medium supplemented with 0 / 0.5 mM trehalose to a final OD_600nm_ ∼0.2. Expression of the reporter was induced with 100 ng/mL anhydrotetracycline (Clontech, #631310) for 6-8 hours. Bacteria were pelleted and lysed with passive lysis buffer provided in the dual-luciferase assay kit (Promega, #E1960), and luminescence was measured by a Fluoroskan Ascent FL luminometer (Thermo Scientific) according to the manufacturer’s instructions, with 1,000 ms as the integration time.

### Statistical analysis

All experiments were performed at least three times independently. The number of repeats is shown in the figure legends. Data were analyzed with unpaired, two-tailed Student’s *t*-test using GraphPad Prism. The results are shown as mean ± standard deviation. The level of significance was set to p < 0.05. *, **, and ***, ns indicated p <0.05, p < 0.01 and p < 0.001, p>0.05 respectively.

## Results

### Mutated *gatA* identified from clinical isolates does not dramatically alter GatCAB enzymatic function

We wished to characterise the glutaminase and amidotransferase activities of mycobacterial GatCAB. We also wanted to compare wild-type enzyme function with two *gatA* mutated enzymes identified in our prior study from clinical isolates of *Mycobacterium tuberculosis*: GatA-K61N and GatA-G444S (9). In that study, we showed that these mutations conferred substantially elevated rates of specific translational error in mycobacteria, which in turn, resulted in enhanced antibiotic tolerance (9). However, the molecular mechanisms by which these mutations resulted in elevated mistranslation rates was not known. The GatA subunit of GatCAB serves as a glutaminase, liberating ammonia from glutamine, which is transferred to GatB via a hydrophilic tunnel where it is used to amidate the misacylated Glu-tRNA^Gln^ or Asp-tRNA^Asn^ substrate (12,13). The glutaminase reaction also results in generation of glutamate from glutamine. Using a commercial luminometric assay for glutamate, we measured the glutaminase activity (Fig. S1) of the wild-type and two mutant enzymes in the presence of Asp-tRNA^Asn^ (Table 1). Although there were some observable differences between the *K*_M_ of the wild-type and in particular the GatCAB-K61N enzyme, the *k*_*cat*_/*K*_M_ of the three enzymes were broadly similar (Table 1). Next, we measured the amidotransferase activity of the three enzymes using Asp-tRNA^Asn^ as a substrate using a modified ^32^P/P1 nuclease assay followed by thin-layer chromatography (27,28) and Fig. S2. Again, the enzymatic activity of all three enzymes were similar, with the two mutant enzymes actually showing slight increases (∼25%) in *k*_*cat*_/*K*_M_ compared with the wild-type enzyme (Table 2). Together, these data suggest that the elevated mistranslation rates observed in mycobacteria harbouring these clinically-derived mutations cannot be explained by decreased enzymatic function due to these mutations.

**Table 1.** Kinetic data for WT, G444S and K61N *M. tuberculosis* GatCAB, glutaminase activity.

| Enzyme | $K_M(\text{Gln, mM})$ | $k_{\text{cat}} (\text{s}^{-1})$ | $k_{\text{cat}}/K_M(\text{Gln, s}^{-1} / \text{mM})$ |
| --- | --- | --- | --- |
| WT | $0.13 \pm 0.02$ | $0.95 \pm 0.04$ | 7.31 |
| G444S | $0.18 \pm 0.03$ | $0.90 \pm 0.04$ | 5.00 |
| K61N | $0.24 \pm 0.04$ | $1.53 \pm 0.08$ | 6.38 |

**Table 2.** Kinetic data for WT, G444S and K61N *M. tuberculosis* GatCAB, amidotransferase activity.

| Enzyme | $K_M$ (Asp-tRNA<sup>Asn</sup>, $\mu\text{M}$) | $k_{\text{cat}}$ ($\text{s}^{-1}$) | $k_{\text{cat}}/K_M$ (Asp-tRNA<sup>Asn</sup>, $\text{s}^{-1} / \mu\text{M}$) |
| --- | --- | --- | --- |
| WT | $0.30 \pm 0.10$ | $0.063 \pm 0.004$ | 0.21 |
| G444S | $0.16 \pm 0.05$ | $0.044 \pm 0.002$ | 0.28 |
| K61N | $0.31 \pm 0.13$ | $0.085 \pm 0.008$ | 0.27 |

Comparison of $K_M$ (with Asp-tRNA<sup>Asn</sup> as substrate), $k_{\text{cat}}$ and $k_{\text{cat}}/K_M$ between GatCAB variants: WT ( $n = 4$ ), G444S ( $n = 4$ ) and K61N ( $n = 3$ ). 1 mM Gln and 4 mM ATP were added with 10 nM enzyme. Values represent means ± SD of independent biological replicates as shown above.

### GatCAB stability is compromised by clinical mutations

Having demonstrated that loss of enzymatic activity does not explain the partial loss of function observed in bacteria with these two mutations, we proceeded to test another aspect of GatCAB function, that of heterotrimer stability. In our earlier study, we had performed an *in vitro* forward genetic selection and screen for high mistranslation mutants and identified several mutations in *gatA* (9). Although the mutations were in *gatA*, (wild-type) GatB protein stability was also lower in a pulse-chase experiment, indirectly implicating heterotrimer stability in those *in vitro* selected mutants (9). We tested two aspects of enzyme stability using thermal ramp assays: conformational stability, determined by measuring complex unfolding through intrinsic fluorescence, denoted Tm1 (Fig. S3A); and colloidal stability by detecting particle aggregation via static light scattering, denoted Tagg266 (Fig. S3B). Both mutant enzymes showed lower colloidal stability with stability of WT>GatCAB-K61N>GatCAB-G444S. With regards to conformational stability, only the GatCAB-G444S was significantly less stable than the other two enzymes (Table 3).

**Table 3.** Thermostability of GatCAB variants (Tm1 and Tagg _266_)

| Enzyme | Tm1 (°C) |  | Tagg<sub>266</sub> (°C) |  |
| --- | --- | --- | --- | --- |
|  | Mean ± SD | p-value | Mean ± SD | p-value |
| WT | 47.2 ± 0.25 | N/A | 46.8 ± 0.28 | N/A |
| G444S | 40.9 ± 0.25 | < 0.001 | 44.1 ± 0.02 | < 0.001 |
| K61N | 48.0 ± 0.44 | 0.06 | 44.7 ± 0.17 | < 0.001 |

Comparison of Tm1 and Tagg<sub>266</sub> between GatCAB variants: WT ( $n = 3$ ), G444S ( $n = 3$ ) and K61N ( $n = 3$ ). Samples were tested at 1.7 mg/mL. p-values show comparison by Student's $t$ -test to WT GatCAB.

### Mycobacterial ND-AspRS and GatCAB form a stable asparagine-transamidosome in a tRNA^Asn^-dependent manner

The transamidosome is a ternary complex of GatCAB, non-discriminatory synthetase and tRNA. Prior studies suggest that several organisms form relatively stable asparaginyl-transamidosomes (17-20,28), however such a complex has never been characterized for a mycobacterial system. We used size-exclusion chromatography to determine whether *M. tuberculosis* also formed a stable asparaginyl-transamidosome (Asn-transamidosome). ND-AspRS, tRNA^Asn^ and GatCAB were co-eluted from a size-exclusion column isolated with a shorter elution time than any of the individual components (Fig. 1A, C). Native PAGE analysis of the individual peaks confirmed a molecular mass shift of the tRNA-containing fraction, in keeping with Asn-transamidosome assembly (Fig. 1B). All three components were necessary for transamidosome formation, since there did not appear to be complex formation between GatCAB and ND-AspRS alone (Fig. S4). These two enzymes eluted with similar retention times as each other, likely reflecting similar masses of GatCAB (117 kDa) and the natural dimeric state (29) of ND-AspRS (130 kDa) and with no shift observed when combined. Both GatCAB-K61N and GatCAB-G444S were also able to form Asn-transamidosomes (Fig. 1D-G).

**Figure 1.**
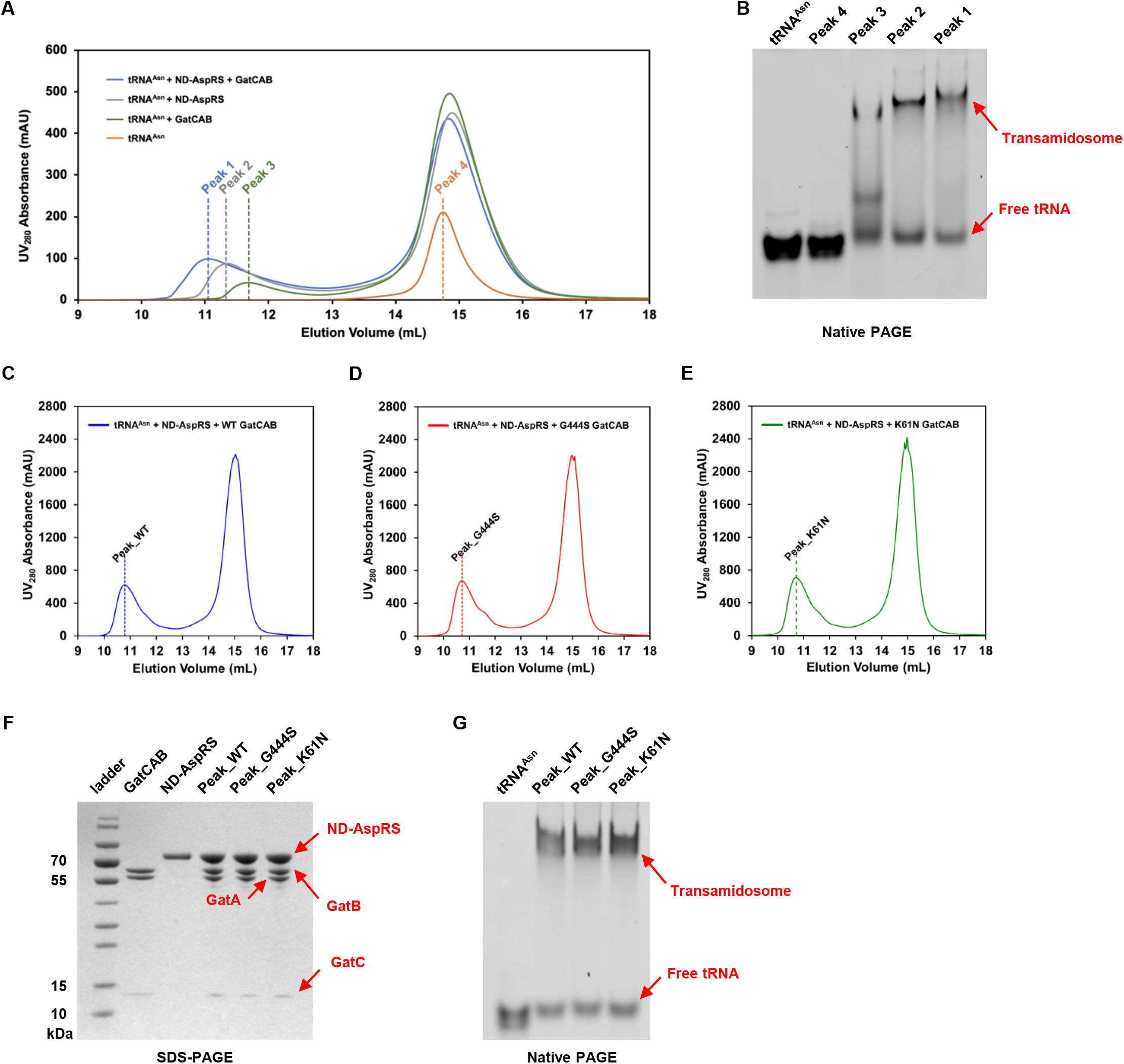
Formation, verification and characterization of the *M. tuberculosis* Asn-transamidosome. **(A)** Size exclusion chromatography of tRNA^Asn^, ND-AspRS and WT GatCAB, confirming formation of the Asn-transamidosome as shown by earlier elution peak. **(B)** Native PAGE analysis of peaks 1-4 from (A), stained with SYBR Green II. Size-exclusion profiles of Asn-transamidosomes with **(C)** WT GatCAB, **(D)** GatCAB-G444S and **(E)** GatCAB-K61N, as well as tRNA^Asn^, ND-AspRS. **(F)** SDS-PAGE analysis of the peaks illustrated in (C-E) alongside WT GatCAB and ND-AspRS. **(G)** Native PAGE of the peaks illustrated in (C-E) alongside tRNA^Asn^ and stained with SYBR Green II.

### The GatCAB-G444S-transamidosome is relatively unstable

To better understand the impact of the GatCAB-K61N and GatCAB-G444S mutations on transamidosome function, we modelled the structure of the mycobacterial Asn-transamidosome based on the crystal structure of the *Pseudomonas aeruginosa* Asn-transamidosome (20). Compared with the wild-type enzyme, the substitution of a serine for the glycine at position 444 in mycobacterial GatA resulted in a predicted steric clash with a proline in the GatC subunit (Fig. 2). No such steric hindrance was observed for GatCAB-K61N (not shown). In keeping with these models, when we measured the stability of the wild-type and mutant enzyme Asn-transamidosomes by thermal ramp assay, colloidal stability was significantly compromised for GatCAB-G444S but not GatCAB-K61N (Table 4).

**Table 4.** Thermostability of Asn-transamidosome variants (Tm1 and Tagg _266_)

| Asn-transamidosome<br>Complex | Tm1 (°C) |  | Tagg<sub>266</sub> (°C) |  |
| --- | --- | --- | --- | --- |
|  | Mean ± SD | p-value | Mean ± SD | p-value |
| WT | 42.0 ± 0.56 | N/A | 43.6 ± 0.44 | N/A |
| G444S | 41.2 ± 1.17 | 0.16 | 43.0 ± 0.44 | 0.03 |
| K61N | 41.7 ± 0.97 | 0.51 | 43.7 ± 0.21 | 0.44 |

Comparison of Tm1 and Tagg<sub>266</sub> between transamidosome variants: WT ( $n = 7$ ), G444S ( $n = 6$ ) and K61N ( $n = 7$ ). Samples were tested at 0.5 mg/mL. p-values show comparison by Student's $t$ -test to WT transamidosome.

**Figure 2.**
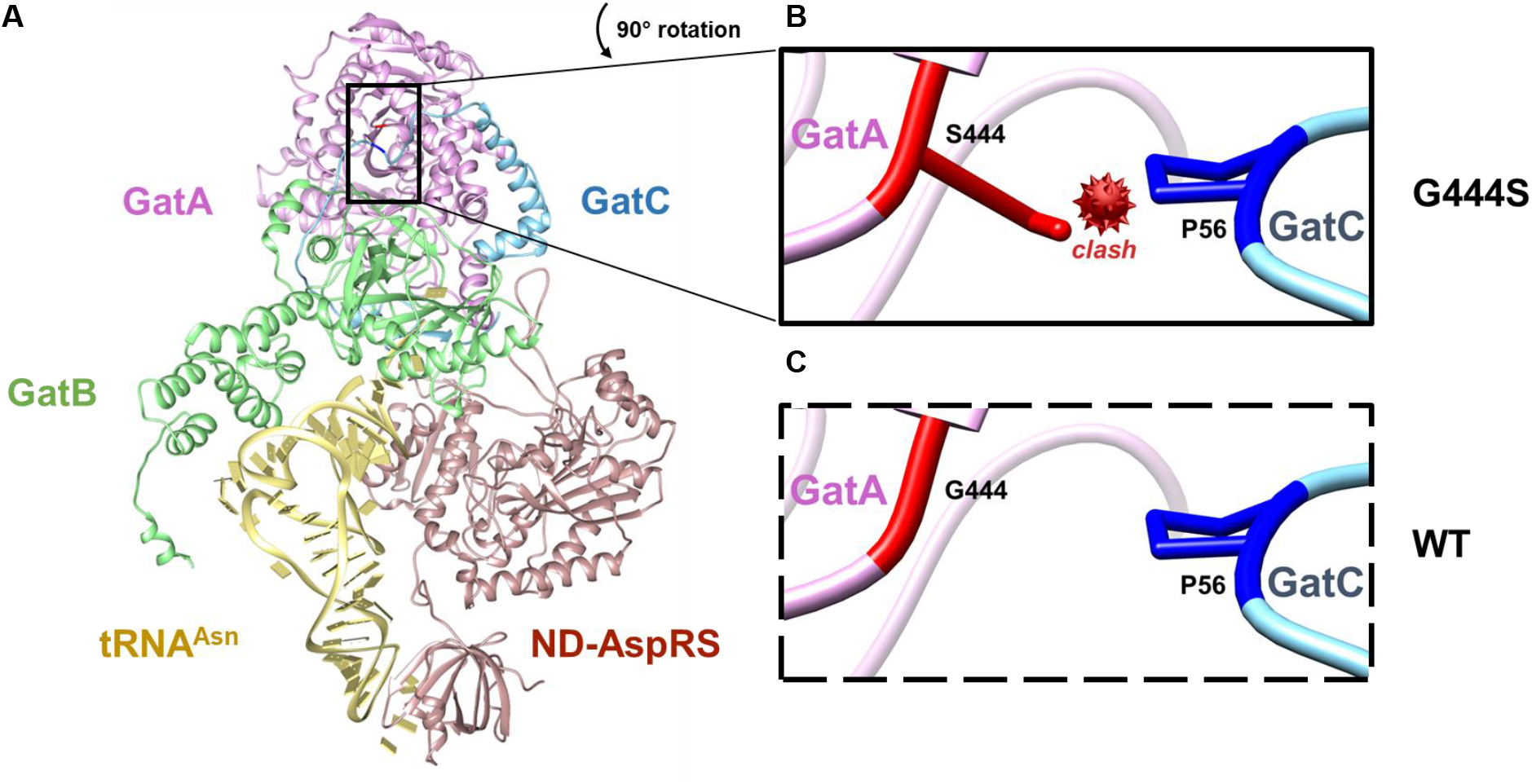
Structural modelling of the *M. tuberculosis* Asn-transamidosome explains complex instability due to GatA-G444S. **(A)** Predicted structural model of the Asn-transamidosome. *M. tuberculosis* GatA, GatB, GatC and ND-AspRS are coloured magenta, green, cyan and brown respectively. tRNA^Asn^ (yellow) docked onto the structure is from the *P. aeruginosa* asparagine transamidosome (PDB ID:4WJ3). The site of the 444 residue of GatA is shown in the boxes **(B – GatA-G444S, C – GatA-WT)**: G444 and S444 are coloured red, while P56 from GatC is blue. The box shows a magnified and 90° counterclockwise rotated view of the mutation site.

### Stabilising GatCAB and the Asn-transamidosome with trehalose increases specific translational fidelity

Our biochemical and biophysical data suggested that *gatA* mutations that impact translational fidelity act via destabilisation of the GatCAB heterotrimer or Asn-transamidosome or both. To further explore this model, we decided to attempt to stabilise the complexes with the sugar trehalose, which has been previously demonstrated to have a buffering effect on potentially deleterious or destabilising mutations (30,31). We repeated the thermostability assays in the absence or presence of trehalose. Trehalose significantly increased the stability of both GatCAB-K61N and GatCAB-G444S heterotrimers (Fig. 3A, B) as well as the corresponding Asn-transamidosomes (Fig. 3C, D). Intriguingly, trehalose also increased the stability of the wild-type GatCAB heterotrimer and Asn-transamidosome (Fig. 3A-D).

**Figure 3.**
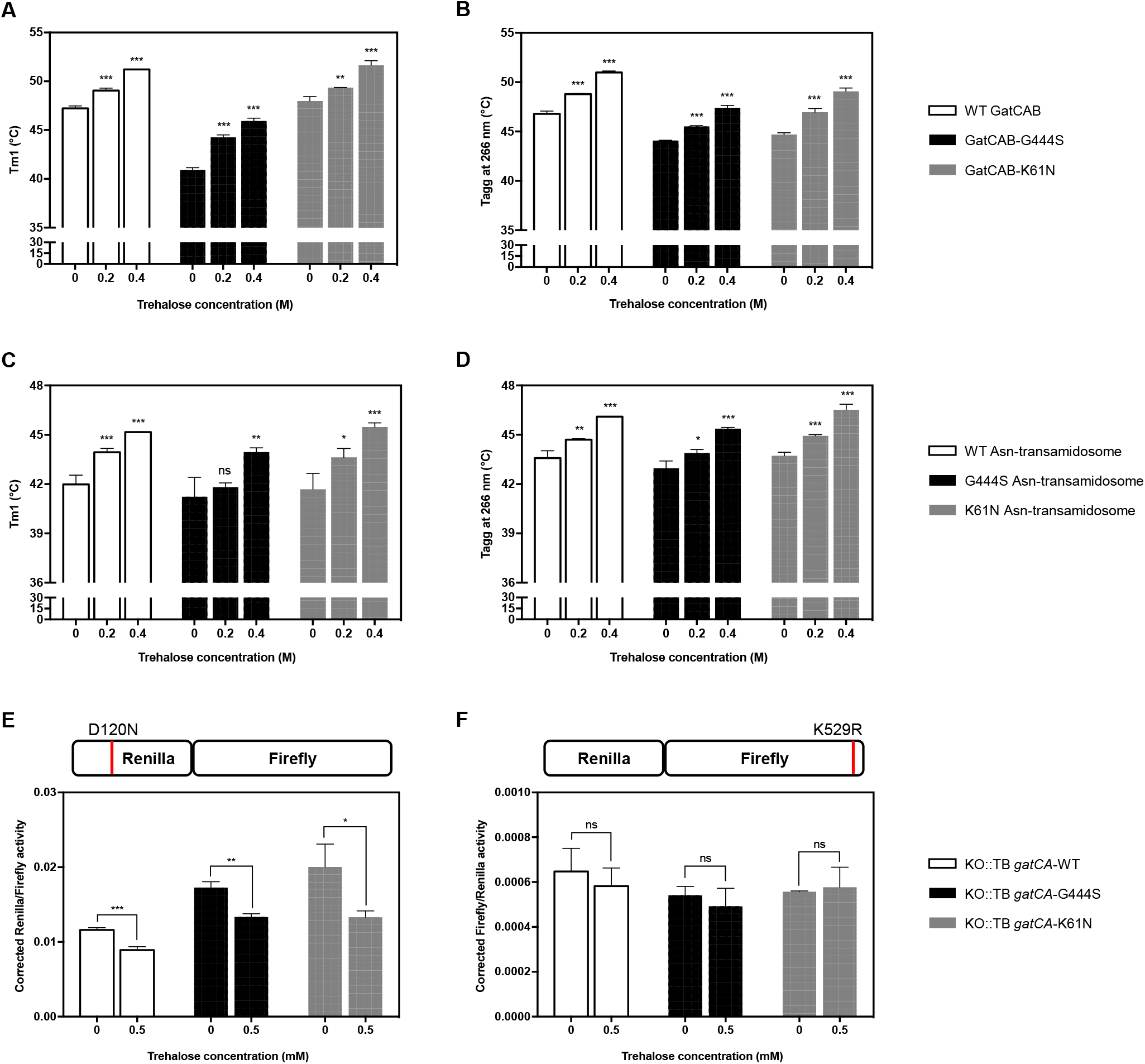
Stabilising GatCAB and the Asn-transamidosome increases specific mycobacterial translational fidelity. *In vitro* thermostability assay of WT and mutant GatCAB **(A – Tm1, B – Tagg**_**266**_) and Asn-transamidosomes **(C – Tm1, D – Tagg**_**266**_) with and without trehalose at indicated concentrations, n ≥3 biological replicates. Mistranslation rates, expressed as corrected Renilla/Firefly **(E)** and corrected Firefly/ Renilla **(F)** of *M. smegmatis* expressing *M. tuberculosis* wild-type or mutated *gatCA* in cultures containing trehalose at indicated concentrations (see Methods). The cartoon above each bar chart depicts the type of error measured (pre-ribosomal error due to the indirect pathway – **E**, and ribosomal decoding errors, **F**). Error bars represent the standard deviation of three independent cultures. Asterisks in each case represent level of significance as determined by Student’s t-test compared with the no trehalose group. ns p> 0.05, * p<0.05, ** p<0.01, *** p<0.001.

To translate these findings to an *in vivo* setting, we decided to test the impact of GatCAB and transamidosome stabilization by trehalose on translational fidelity in mycobacteria. For pragmatic reasons including the lack of a luminometer in the biosafety level-3 facility, we tested mistranslation rates in the non-pathogenic *Mycobacterium smegmatis*, in which the native *gatCA* locus had been deleted and replaced elsewhere on the chromosome with wild-type or mutant *gatCA* from *M. tuberculosis* (9). Using a set of dual-luciferase reporters (7,32), we decided to measure two different forms of translational error representing two distinct pathways. To quantify errors generated by the pre-ribosomal indirect tRNA aminoacylation pathway we measured mistranslation of aspartate for asparagine using a reporter in which a critical aspartate residue in Renilla luciferase was mutated to asparagine (25). We also measured ribosomal decoding errors using the same Renilla-Firefly luciferase system, but with a mutation of a critical lysine in Firefly luciferase (26,32,33). We reasoned that if trehalose specifically stabilised GatCAB and the Asn-transamidosome, only the pre-ribosomal error rate should be lowered. Indeed, asparagine to aspartate mistranslation rates were significantly lower with trehalose not only in the strains expressing mutant *gatA*, but also in the strain expressing wild-type *M. tuberculosis gatA* (Fig. 3E). By contrast, there was no significant impact of trehalose supplementation on rates of ribosomal decoding errors in any of the strains, suggesting that trehalose was not influencing translational fidelity or reporter function in a general manner (Fig. 3F). These data suggest that stabilisation of wild-type, as well as mutant GatCAB and/or the Asn-transamidosome increases translational fidelity of the indirect tRNA aminoacylation pathway and suggests this may be a physiologically relevant mechanism by which mistranslation rates are tuned in response to environmental triggers and stressors (7,9).

## Discussion

In this study, we chose to focus on two mutations in *gatA* identified from tuberculosis patient samples and previously confirmed to cause increased rates of specific mistranslation and antibiotic tolerance (9). Both the GatCAB-K61N and GatCAB-G444S mutations destabilised the heterotrimeric enzyme as measured by thermal ramp assays, but only the GatCAB-G444S-Asn-transamidosome was less stable than wild-type using the same assays, and the relative difference in stability of the transamidosome was smaller than for just GatCAB. There are two possible explanations for these findings. Prior to assay by thermal ramp, the Asn-transamidosomes were first purified by gel-filtration. This would pre-select the samples for stable complex, since highly unstable complexes would not elute as the higher molecular weight peak. Additionally, the assembly of the ternary complex itself may further stabilise (17) and mitigate against the destabilising mutation. By bringing together all three components required for the generation of cognately aminoacylated tRNA^Asn^, the Asn-transamidosome has been proposed to increase fidelity of translation in the indirect pathway (17). Our studies confirm not only that mutations that destabilise the ternary complex also cause increased mistranslation, but that stabilising the complex, in this case with the disaccharide trehalose, improves translational fidelity: demonstrating the validity of the model in living bacteria for the first time.

Although our kinetic studies found that the enzymatic function of the GatCAB-K61N and GatCAB-G444S mutants were broadly similar to the wild-type enzyme, although there were subtle decreases in glutaminase activity. However, glutaminase is not the rate-limiting step in GatCAB function, and the mutant enzymes actually had slightly increased amidotransferase activity. One recent study performed a modified glutaminase assay for mycobacterial GatCA B (34). However, they did not conduct kinetic analyses, and their assay conditions – extremely high enzyme concentrations and incubation over several hours prior to measurements, make comparison with our studies difficult. More generally, our kinetic studies suggest that mycobacterial GatCAB is less efficient than enzymes previously characterized from *Neisseria meningitidis* and *Moraxella catarrhalis*, although more similar to that of *Helicobacter pylori* (19,27,35,36). Relatively inefficient amidotransferase activity by mycobacterial GatCAB may explain the observed high basal mistranslation associated with this pathway in mycobacteria, however it should be noted that the equivalent error rates in the other organisms have not been measured. It may also explain why individual bacilli within an isogenic population of wild-type mycobacteria that have lower expression of (wild-type) *gatCAB* have higher mistranslation rates, suggesting that GatCAB expression is limiting in mycobacteria (9). The activity of *H. pylori* GatCAB was stimulated by the presence of an additional protein, Hp0100 (QueH), which also allowed the formation of a stable, Hp0100-dependent but tRNA-independent Asn-transamidosome (28). It is possible that a functionally equivalent protein exists in mycobacteria, boosting both GatCAB activity and transamidosome stability. However, it should be noted that unlike *Helicobacter* (19,28), the mycobacterial Asn-transamidosome is stable with just the tRNA, synthetase and GatCAB components.

Small molecules targeting enzymatic inhibition of GatCAB have been discovered (37,38), but none have been validated against bacteria. Our work suggests that compounds that inhibit GatCAB could act not just by inhibiting enzyme synthesis, but also by destabilising GatCAB or Asn-transamidosome complex formation. Alternatively, compounds that hyper-stabilise the complex may decrease mistranslation and consequently reduce ‘adaptive mistranslation’ analogous to the effect of kasugamycin on rifampicin susceptibility (10). Given the presence of the indirect tRNA aminoacylation pathway in the majority of human pathogens, further studies in targeting this pathway may be warranted. Studies of naturally occurring mutations in the genes encoding GatCAB can inform complex function. Mutations in all three components of human, mitochondrial GatCAB have been associated with neonatal lethal cardiomyopathy (39). This may suggest that mammalian mitochondria are less tolerant of partial loss-of-function mutations in GatCAB, but it is noteworthy that there was a clear gradient in toxicity caused by the mutations in tissues, with cardiac muscle severely affected and almost no phenotype in patient fibroblasts (39). By contrast, bacteria seem to better tolerate mutations in *gatCAB*. Penicillin-resistant isolates of *M. catarrhalis* have an insertion of a beta-lactamase gene between *gatA* and *gatB* in the *gatCAB* operon. The insertion alters the C-terminus of GatA, resulting in a subtle change in enzyme activity, but with no apparent loss in bacterial viability (35). Analysis of the two mycobacterial *gatA* mutations presented here suggest a potential mechanism by which mycobacteria may tune translational fidelity in response to environmental cues (7). Having identified that the mutations destabilise GatCAB and the Asn-transamidosome, we sought to stabilise the complexes with trehalose. Trehalose is a naturally occurring glucose disaccharide that is found in bacteria, plants and some invertebrates but absent in mammals. Trehalose is essential for mycobacterial growth (40) and cell-wall component synthesis (41,42), and abundant in mycobacteria, making up 1-3% of dry weight. Trehalose has also been implicated in stress response and buffering of deleterious mutations, acting as a chemical chaperone (30,31,43). We found that trehalose not only stabilised mutant GatCAB but also wild-type GatCAB and the Asn-transamidosome. Importantly, trehalose supplementation of cultures of mycobacterial strains expressing wild-type GatCAB caused a reduction in translational errors specifically for the indirect tRNA aminoacylation pathway. Our work suggests that studying naturally occurring variations in essential members of the translation apparatus can inform mechanisms of protein synthesis regulation.

## Author contributions

YYL and RJC designed research, performed the majority of experiments and analysed data. JYY and YX performed the structural model. TLH performed the initial training of RJC on the amidotransferase assay. BJ conceived of the study, designed experiments and analysed data. YYL and BJ wrote the manuscript with input from RJC, TLH and the other authors.

## Acknowledgements

We would like to acknowledge the Center of Protein Research and Technology at Tsinghua University for use of their UNcle instrument and ÄKTA purifier, and the Center of Biomedical Analysis at Tsinghua University for use of their isotope facilities. We would also like to thank Hong-Wei Su for constructing *Mycobacterium smegmatis* strains with Mtb WT, G444S and K61N GatCAB. This work was funded in part by grants from the Bill & Melinda Gates Foundation (OPP1109789) and funds from Tsinghua University School of Medicine to BJ. BJ is a Wellcome Trust Investigator (207487/C/17/Z).

