## Supplementary Figures for "Clinically relevant mutations of mycobacterial GatCAB inform regulation of translational fidelity"

A

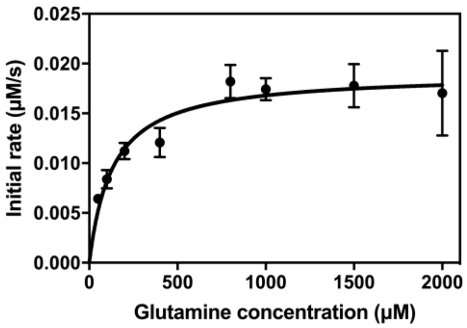


B

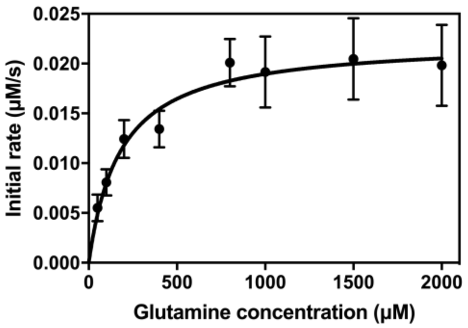


C

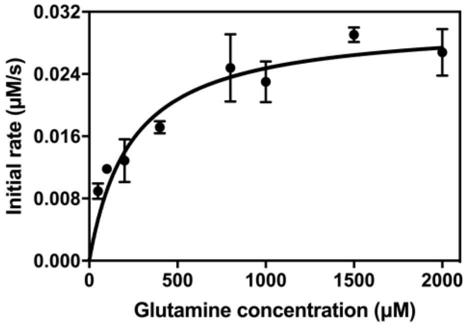


D

| Glutamine concentration (μM) | *n* of different GatCAB groups | | |
|---|---|---|---|
| | WT | G444S | K61N |
| 50 | 4 | 5 | 3 |
| 100 | 4 | 5 | 3 |
| 200 | 4 | 5 | 7 |
| 400 | 5 | 6 | 5 |
| 800 | 4 | 4 | 5 |
| 1000 | 5 | 4 | 3 |
| 1500 | 4 | 4 | 3 |
| 2000 | 4 | 4 | 3 |

**Supplementary Figure 1. Glutaminase assay of WT, G444S and K61N *M. tuberculosis* GatCAB.** Michaelis-Menten curves of **(A)** WT **(B)** G444S and **(C)** K61N GatCAB glutaminase activity. Reactions were carried with 5 μM Asp-tRNA$^{Asn}$, containing ~20 nM GatCAB. GraphPad Prism was used to calculate kinetic parameters by nonlinear regression. Error bars represent the standard deviation of independent biological replicates, the *n* of which are listed in **(D)**.

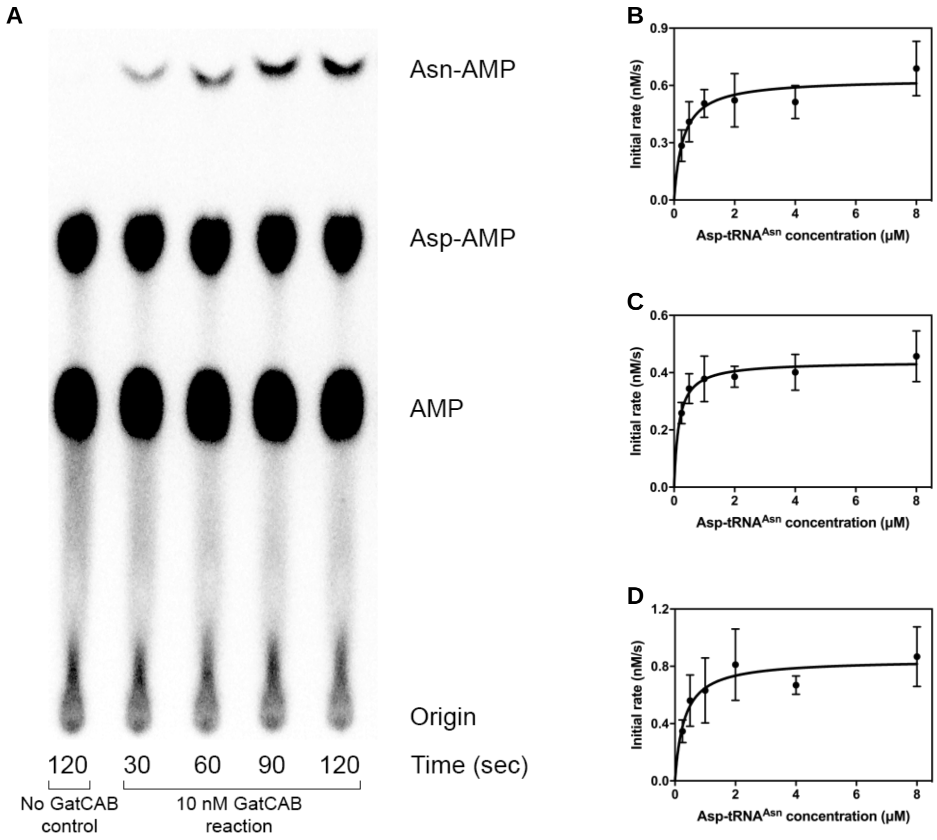


**Supplementary Figure 2. GatCAB amidotransferase assay with $^{32}$P-labeled tRNA.**
**(A)** Representative phosphorimage of the separation of Asn-[α-$^{32}$P]AMP, Asp-[α-$^{32}$P]AMP, [α-$^{32}$P]AMP, by PEI cellulose chromatography. Aliquots (5 μL) of the amidotransferase reaction at 37 °C (10 nM GatCAB from *M. tuberculosis* , 0.5 μM Asp-tRNA$^{Asn}$, 1 mM Gln, 4 mM ATP, 25 mM KCl, 8 mM $MgCl_2$, 40 mM HEPES-KOH pH 7.5) were taken at the time points indicated and quenched / digested at 37 °C with 5 μL of 100 mM sodium citrate pH 4.7, and 0.66 mg/mL of nuclease P1. Digested samples (3 μL) were then spotted onto a 20 × 20 cm PEI-cellulose glass plate, which was developed in 10 mM ammonium chloride, 5% acetic acid for ~2 hours. Aliquot from no GatCAB reaction at 120 seconds was taken as the background control. Michaelis-Menten curves of WT (*n* = 4) **(B)**, G444S (*n* = 4) **(C)** and K61N (*n* = 3) **(D)** GatCAB are shown for their amidotransferase activity. GraphPad Prism was used to calculate kinetic parameters by nonlinear regression. Error bars represent the standard deviation of independent biological replicates.

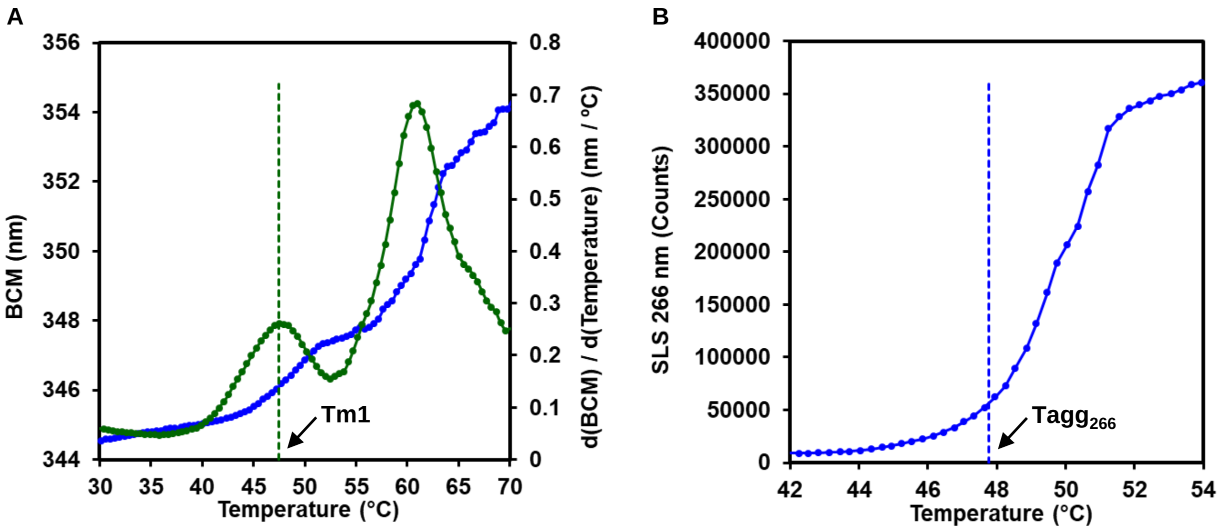


**Supplementary Figure 3. Thermostability assay of *M. tuberculosis* GatCAB.**
The assay was conducted on an UNcle instrument (Unchained Labs) by monitoring thermal melting and aggregation over a temperature range. **(A)** Representative Tm1 characterization of WT GatCAB using intrinsic fluorescence. At each temperature, the barycentric mean (BCM) of the intrinsic fluorescence spectra was measured and plotted against temperature to generate the BCM curve (blue). First-order derivative of the BCM curve was calculated and plotted against temperature (green). The temperature corresponding to the first peak of the first-order derivative curve was defined as Tm1, as shown by the dotted line (green). **(B)** Representative $Tagg_{266}$ characterization of WT GatCAB using static light scattering. At each temperature, the intensity of static light scattering at 266 nm (as represented by SLS 266 nm) was measured and plotted against temperature to generate an aggregation curve. Aggregation onset temperature ($Tagg_{266}$) of each protein was calculated by Uncle Analysis software, as shown by the dotted line.

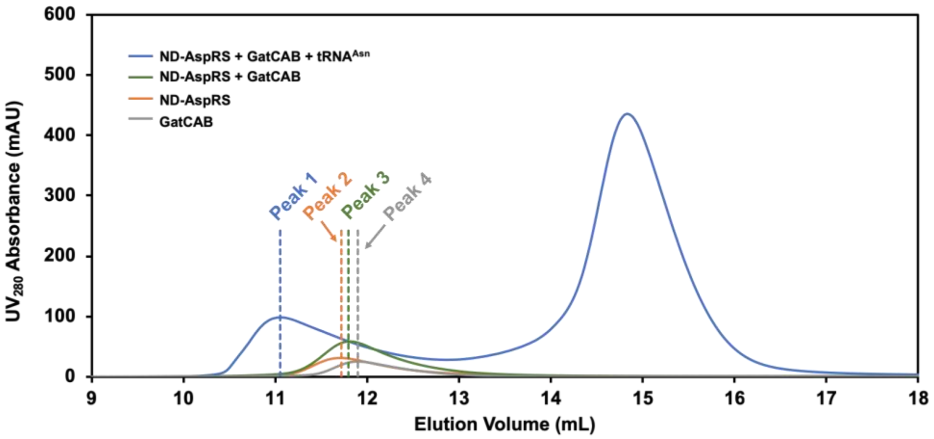


**Supplementary Figure 4. Formation of the *M. tuberculosis* Asn-transamidosome is tRNA-dependent.** Gel filtration conducted with free ND-AspRS, free GatCAB, a mixture of ND-AspRS and GatCAB, and a mixture of ND-AspRS, GatCAB and $tRNA^{Asn}$. 4 µM ND-AspRS, 2 µM WT GatCAB and 2 µM $tRNA^{Asn}$ were used.
