## Supplementary tables for "Clinically relevant mutations of mycobacterial GatCAB inform regulation of translational fidelity"

**Table S1. Bacterial strains used in this study.**

| Name | Descriptions | References or source |
| --- | --- | --- |
| *E. coli*, pET28a-AspS-CHis | Strain for expressing Mtb ND-AspRS, with C-terminal 6×His tag | This study |
| *E. coli*, pETDuet1-NStrep-GatCA-WT-GatB-CHis | Strain for expressing Mtb WT GatCAB, with N-terminal Strep tag II on GatC, and C-terminal 6×His tag on GatB | This study |
| *E. coli*, pETDuet1-NStrep-GatCA-G444S-GatB-CHis | Strain for expressing Mtb G444S GatCAB, with N-terminal Strep tag II on GatC, and C-terminal 6×His tag on GatB | This study |
| *E. coli*, pETDuet1-NStrep-GatCA-K61N-GatB-CHis | Strain for expressing Mtb K61N GatCAB, with N-terminal Strep tag II on GatC, and C-terminal 6×His tag on GatB | This study |
| *E. coli*, pTrc99a-T7-tRNA^Asn^ | Strain for transcripting Mtb tRNA^Asn^ under T7 promoter | This study |
| *M. smegmatis* KO::*gatCA*-WT, pTet-Ren-FF | Reporter strain where the WT *gatCA* gene is replaced by Mtb WT *gatCA* gene, transformed with Renilla-Firefly dual luciferase construct | (9) |
| *M. smegmatis* KO::*gatCA*-WT, pTet-Ren-D120N-FF | Reporter strain where the WT *gatCA* gene is replaced by Mtb WT *gatCA* gene, transformed with mutated Renilla-Firefly dual luciferase construct: measures Asn-to-Asp mistranslation | (9) |
| *M. smegmatis* KO::*gatCA*-WT, pTet-Ren-FF-K529R | Reporter strain where the WT *gatCA* gene is replaced by Mtb WT *gatCA* gene, transformed with mutated Renilla-Firefly dual luciferase construct: measures Arg-to-Lys mistranslation | (9, 26) |
| *M. smegmatis* KO::*gatCA*-G444S, pTet-Ren-FF | Reporter strain where the WT *gatCA* gene is replaced by Mtb G444S *gatCA* gene, transformed with Renilla-Firefly dual luciferase construct | (9) |
| *M. smegmatis* KO::*gatCA*-G444S, pTet-Ren-D120N-FF | Reporter strain where the WT *gatCA* gene is replaced by Mtb G444S *gatCA* gene, transformed with mutated Renilla-Firefly dual luciferase construct: measures Asn-to-Asp mistranslation | (9) |
| *M. smegmatis* KO::*gatCA*-G444S, pTet-Ren-FF-K529R | Reporter strain where the WT *gatCA* gene is replaced by Mtb G444S *gatCA* gene, transformed with mutated Renilla-Firefly dual luciferase construct: measures Arg-to-Lys mistranslation | (9, 26) |
| *M. smegmatis* KO::*gatCA*-K61N, pTet-Ren-FF | Reporter strain where the WT *gatCA* gene is replaced by Mtb K61N *gatCA* gene, transformed with Renilla-Firefly dual luciferase construct | (9) |
| *M. smegmatis* KO::*gatCA*-K61N, pTet-Ren-D120N-FF | Reporter strain where the WT *gatCA* gene is replaced by Mtb K61N *gatCA* gene, transformed with mutated Renilla-Firefly dual luciferase construct: measures Asn-to-Asp mistranslation | (9) |
| *M. smegmatis* KO::*gatCA*-K61N, pTet-Ren-FF-K529R | Reporter strain where the WT *gatCA* gene is replaced by Mtb K61N *gatCA* gene, transformed with mutated Renilla-Firefly dual luciferase construct: measures Arg-to-Lys mistranslation | (9, 26) |

**Table S2. Primers used in this study.**

| Name | Sequence (5’ to 3’) | Description |
| --- | --- | --- |
| *aspS*-F | TCTAGAAAGGAGATATACC**ATG**TTTGTGCTGCGCAGCCACG (*Xba*I underlined) | Used to amplify Mtb *aspS*-CHis (Rv2572c, encoding Mtb ND-AspRS, start codon changed to ATG) |
| *aspS*-R | AAGCTTTCATCAGTGGTGGTGGTGGTGGTGTGCCTGCTGGACCCGCTTG (*Hind*III and His_6_ tag underlined respectively) |  |
| NStrep-*gatCA*-F1 | GAAGGAGATATACC**ATG**TGGAGCCACCCGCAGTTCGAAAAGTCCCAGATCTCCCGCGAC (Strep tag II underlined) | First-step PCR to amplify Mtb NStrep-*gatCA* (start codon of *gatC* changed to ATG) |
| NStrep-*gatCA*-R1 | TTCCCCTATAGTGAGTCGTATTAGGTACCGAATTCTCATCAAATGGCGCTCAGTAGCGGG |  |
| NStrep-*gatCA*-F2 | CTTTAAGAAGGAGATATACCATGTG | Second-step PCR to amplify Mtb NStrep-*gatCA* |
| NStrep-*gatCA*-R2 | TTCCCCTATAGTGAGTCGTATTAG |  |
| *gatB*-CHis-F1 | CATCTTAGTATATTAGTTAAGTATAAGAAGGAGATATACC**ATG**ACTGTTGCTGCCGGGGCAG | First-step PCR to amplify Mtb *gatB*-CHis |
| *gatB*-CHis-R1 | TCATCAATGATGATGATGATGATGACCCTGCCCGCAGGCCTC (His_6_ tag underlined) |  |
| *gatB*-CHis-F2 | GACTCACTATAGGGGAATTGTGAGCGGATAACAATTCCCCATCTTAGTATATTAG | Second-step PCR to amplify Mtb *gatB*-Chis |
| *gatB*-CHis-R2 | GCAGCAGCCTAGGTTAACTCGAGTCATCAATGATGATGATGATGATGACCCTG |  |
| *gatA*-ATG-F | GGGGGATGAACA**ATG**ACGGACATCATCCGAT | Changing start codon of WT *gatA* to ATG using site-directed mutagenesis (SDM) |
| *gatA*-ATG-R | ATCGGATGATGTCCGTCATTGTTCATCCCCC |  |
| *gatA*-G444S-F | CTGCCGCTGAACTTGGCC**AGC**CACTGCGGCATGTCTG | Construction of pETDuet1-NStrep-GatCA-G444S-GatB-CHis using SDM |
| *gatA*-G444S-R | CAGACATGCCGCAGTGGCTGGCCAAGTTCAGCGGCAG |  |
| *gatA*-K61N-F | GCGGCCGCCATCGAC**AAT**CAGGTGGCCGCTGGAGAACC | Construction of pETDuet1-NStrep-GatCA-K61N-GatB-CHis using SDM |
| *gatA*-K61N-R | GGTTCTCCAGCGGCCACCTGATTGTCGATGGCGGCCGC |  |
| T7-tRNA^Asn^-F | CGGAATTC*taatacgactcactata*TCCCCTGTAGCTCAATTGGCAGAGCGTTCGGCTGTTAACCGAAG (*EcoR*I underlined; T7 promoter sequence in italics) | Construstion of pTrc99a-T7-tRNA^Asn^ |
| T7-tRNA^Asn^-R | CGGGATCCTGGCTCCCCCGGGAGGACTCGAACCTCCAACCCTTCGGTTAACAGCCGAACGCTCTG (*BamH*I underlined) |  |
